# Error-correcting DNA barcodes for high-throughput sequencing

**DOI:** 10.1101/315002

**Authors:** John A. Hawkins, Stephen K. Jones, Ilya J. Finkelstein, William H. Press

**Affiliations:** Institute for Computational Engineering and Science, The University of Texas at Austin, Austin, TX 78712, USA; Department of Molecular Biosciences, The University of Texas at Austin, Austin, TX 78712, USA; Institute for Cellular and Molecular Biology, The University of Texas at Austin, Austin, TX 78712, USA; Center for Systems and Synthetic Biology, The University of Texas at Austin, Austin, TX 78712, USA; Department of Integrative Biology, The University of Texas at Austin, Austin, TX 78712, USA

**Keywords:** error-correcting codes, next-generation sequencing, DNA barcodes, DNA library, massively parallel synthesis

## Abstract

Many large-scale high-throughput experiments use DNA barcodes—short DNA sequences prepended to DNA libraries—for identification of individuals in pooled biomolecule populations. However, DNA synthesis and sequencing errors confound the correct interpretation of observed barcodes and can lead to significant data loss or spurious results. Widely-used error-correcting codes borrowed from computer science (e.g., Hamming and Levenshtein codes) do not properly account for insertions and deletions in DNA barcodes, even though deletions are the most common type of synthesis error. Here, we present and experimentally validate FREE (Filled/truncated Right End Edit) barcodes, which correct substitution, insertion, and deletion errors, even when these errors alter the barcode length. FREE barcodes are designed with experimental considerations in mind, including balanced GC content, minimal homopolymer runs, and reduced internal hairpin propensity. We generate and include lists of barcodes with different lengths and error-correction levels that may be useful in diverse high-throughput applications, including >10^6^ single-error correcting 16-mers that strike a balance between decoding accuracy, barcode length, and library size. Moreover, concatenating two or more FREE codes into a single barcode increases the available barcode space combinatorially, generating lists with > 10^15^ error-correcting barcodes. The included software for creating barcode libraries and decoding sequenced barcodes is efficient and designed to be user-friendly for the general biology community.

**SIGNIFICANCE STATEMENT:** Modern high-throughput biological assays study pooled populations of individual members by labeling each member with a unique DNA sequence called a “barcode.” DNA barcodes are frequently corrupted by DNA synthesis and sequencing errors, leading to significant data loss and incorrect data interpretation. Here, we describe a novel error-correction strategy to improve the efficiency and statistical power of DNA barcodes. To our knowledge, this is the first report of an error-correcting method that accurately handles insertions and deletions in DNA barcodes, the most common type of error encountered during DNA synthesis and sequencing, resulting in order-of-magnitude increases in accuracy, efficiency, and signal-to-noise. The accompanying software package makes deployment of these barcodes effortless for the broader experimental scientist community.

## INTRODUCTION

Many modern large-scale biology experiments use high-throughput DNA sequencing to study the behavior of individual biomolecules in pooled populations. These experiments encode the identity of individual members via DNA barcodes—short, unique DNA sequences that are coupled to each member in the population (Fig. 1a). DNA barcode-based identification is central to such diverse applications as single-cell genome and RNA sequencing^1–7^, gene synthesis^8,9^, high-throughput antibody screens^10,11^, and drug discovery^12,13^. Such experiments have been enabled by recent breakthroughs in massively-parallel, pooled DNA synthesis^14,15^. For example, a recent study used DNA barcodes to discover small molecule inhibitors of enzymes by screening ~10^8^ small molecules. Each small molecule was attached to a unique set of three DNA barcodes. The highest affinity ligands were enriched via multiple rounds of selection and then identified via high-throughput sequencing of the attached barcodes^16^. The rapid growth of such methodologies in all areas of biomedicine requires the development of large pools (>10^6^ members) of unique DNA barcodes to identify individual members (e.g., cells, proteins, drugs) in heterogeneous ensembles.

**Figure 1:**
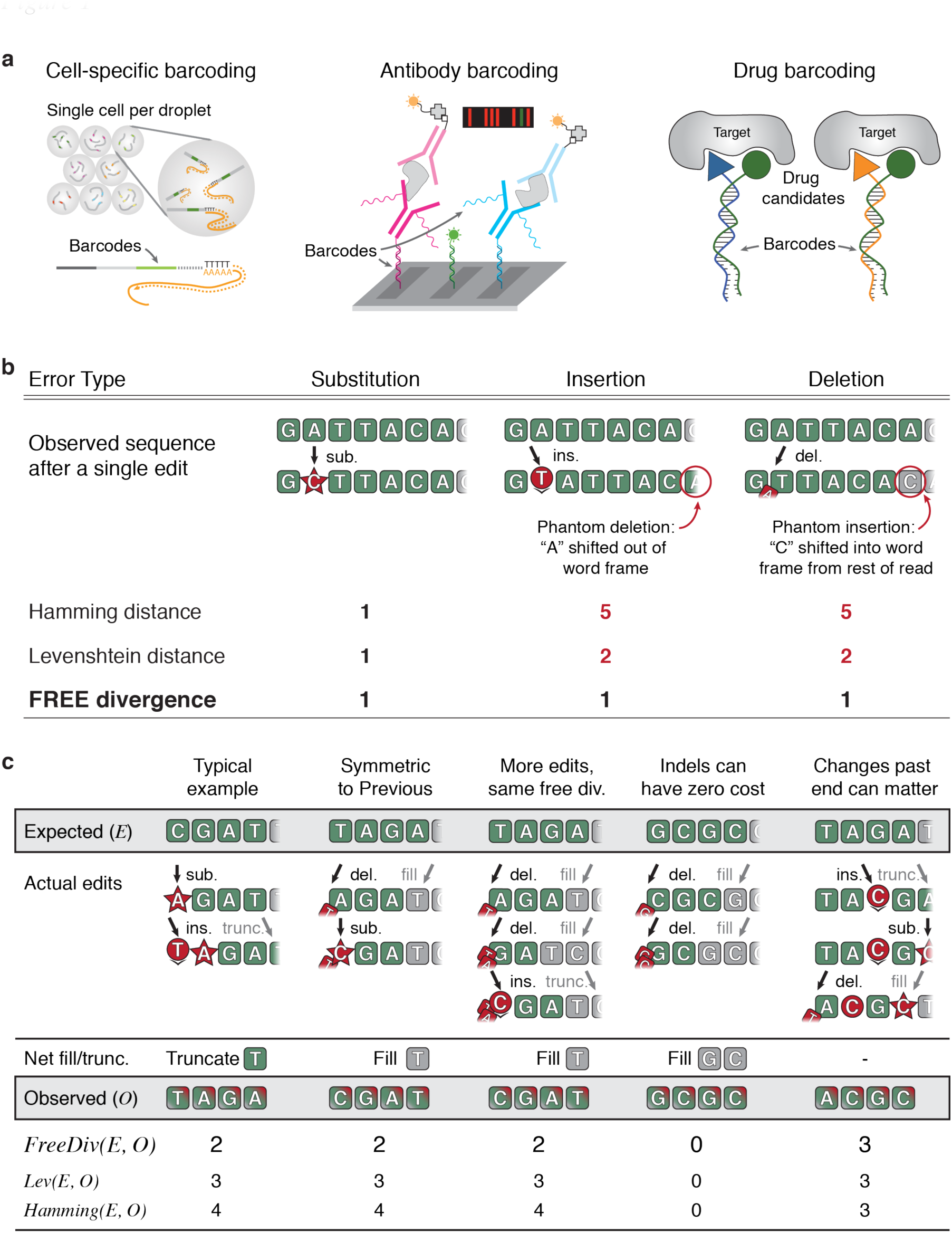
Applications and error-correction strategies of DNA barcodes. **a.** Illustrative examples of high-throughput sequencing assays that require large lists of error-correcting DNA barcodes. Barcodes are used to identify individual cells or molecules in pooled libraries (Klein, 2015; Fan, 2008; Melkko, 2004). **b.** Current strategies to correct synthesis and sequencing errors in DNA barcodes are confounded by insertions and deletions. Hamming distance can only handle substitutions. Levenshtein distance is confounded by the fact that barcodes are prepended to other sequences of interest. Indels thus produce phantom Levenshtein distance errors when bases from the remaining DNA molecule shift into or out of the barcode window. **c.** Examples of FREE divergence (this work) given the actual edit history. Levenshtein and Hamming distances are also shown for comparison. A substitution and insertion are correctly attributed as 2 edits by FREE divergence (first column). FREE divergence is a symmetric function, i.e., *FreeDiv(E, O) = FreeDiv(O, E)* (first and second columns). Different actual edit paths can result in the same observed sequence (second and third columns). Indels can have zero cost, particularly near the end of the barcode where they can occasionally be undone by fill or truncation (fourth column). Edits past the barcode end can matter since the fill/truncation step happens only upon observation (fifth column).

Every assay with DNA barcodes is subject to errors introduced during DNA synthesis and sequencing. These errors decrease experimental power and accuracy by confounding the identity of individual biomolecules in the population. The most common DNA synthesis error is a single-base deletion (Results). This is particularly challenging to decode because it causes a frameshift in all downstream sequencing. Substitutions and insertion errors are also common during massively-parallel pooled oligonucleotide synthesis (Results). Our own experimental results are consistent with manufacturer-advertised error rates of up to 1 per 200 nucleotides (nt)^17^. For 20 base pair (bp) long barcodes with no error correction, this translates to a best-case scenario of 10% data lost or, worse, incorrectly interpreted. Next-generation sequencing also has error rates between 10^−3^ and 10^−4^. This alone represents errors in approximately 1% of our example 20 bp barcodes, which can be limiting for detection of rare events. These errors can be overcome through the use of error-correcting DNA barcodes—DNA sequences that can correctly identify the underlying individuals in a pooled experiment even in the presence of sequencing and synthesis errors.

Error-correcting barcodes must efficiently detect and correct all DNA sequencing and synthesis errors. Many current DNA barcode strategies repurpose error-correcting codes developed for computers^18,19^, such as Hamming or Reed-Solomon codes, to DNA applications^20,21^. Hamming distance, i.e., the number of substitutions between two sequences of equal length, is possibly the most used due to its simplicity. However, nearly all well-studied error-correcting codes developed in computer science—including the widely-used Hamming codes—were not designed to handle deletions and insertions, which are the most common errors in DNA synthesis. Such codes are generally used to only detect errors without correcting them, but even then there is a possibility that a single error (e.g., deletion) can convert one barcode into another. Levenshtein codes, also known as edit codes, can theoretically account for all three types of common error: substitutions, insertions, and deletions, but only when the corrupted length of each barcode after errors is known^22,23^. This is a critical limitation in real-world DNA barcode applications because errors can change the barcode length unpredictably, which leads to erroneous decoding of Levenshtein-based barcodes in the context of a longer read (Fig. 1b). As a workaround, Levenshtein codes can be used at twice the level of error correction as desired for a given application, for example using a 2 error-correcting code when a 1-error correcting code is desired, but this is inefficient and significantly decreases the number of valid barcodes for a given oligonucleotide length. In sum, existing DNA barcode strategies are unable to efficiently detect and decode real-world errors encountered during DNA synthesis and sequencing.

Here, we develop and experimentally validate error-correcting Filled/truncated Right End Edit (FREE) barcodes. FREE barcodes can correct substitutions, insertions, and deletions even when the edited length of the barcode is unknown. These barcodes are designed with experimental considerations in mind, including balanced GC content, minimal homopolymer runs, and no self-complementarity of more than two bases to reduce internal hairpin propensity. We generate and include lists of barcodes with different lengths and error-correction levels that may be broadly useful in diverse high-throughput applications. For each barcode set, we calculate hairpin melting temperatures which can be used to select subsets of barcodes to match experimental conditions. Our largest barcode list includes >10^6^ unique error-correcting barcodes usable in a single experiment. Moreover, appending two or more barcodes together combinatorially increases the total barcode set, producing >10^9^-10^12^ unique error-correcting DNA barcodes. The included software for creating new barcode libraries and decoding/error-correcting observed barcodes is fast and efficient, decoding >120,000 barcodes per second with a single processor, and is designed to be user friendly for a broad biologist community.

## RESULTS

### Overview of <u>F</u>illed/truncated <u>R</u>ight <u>E</u>nd <u>E</u>dit (FREE) Divergence Codes

After DNA synthesis and sequencing, a barcode of length *n* can be altered, and is not guaranteed to end after exactly *n* bases. Our goal is to design barcodes that can be unambiguously identified from the first *n* bases of the sequenced read. To begin, we define a *filled/truncated right-end m-edit*, hereafter written “FRE m-edit,” of a DNA sequence of length *n* to be the result of any *m* edits—substitutions (*sub*), insertions (*ins*), or deletions (*del*)—followed by truncating or filling with any random bases on the right (as from the unknown downstream read) as necessary to return to original length *n* (Fig. 1b). For any two DNA sequences *X* and *Y* of the same length, we define the *Filled/truncated Right End Edit (FREE) Divergence* between *X* and *Y,* written *FreeDiv(X, Y),* to be the minimum *m* such that *Y* is a FRE *m*-edit of *X*.

Figure 1c shows a typical example of how FREE divergence captures the actual number of barcode edits in the context of a longer read. An insertion has caused the final T to move out of the barcode window, but FREE divergence correctly accounts for its loss. FREE divergence is a symmetric function, i.e. *FreeDiv(X, Y) = FreeDiv(Y, X)* (Fig. 1c). This is because reversing the edits and reversing the right-end fill or truncation step moves one from *Y* back to *X* in the same minimum number of steps (Supplemental Materials). FREE divergence is defined as the minimum number of steps between the expected and observed barcode, but it is possible to accomplish the same transformation with more edits, for example via the identity *ins-del = sub* (Fig. 1c). Also, insertions and/or deletions (indels) near the end of the sequence can result in a FREE divergence of zero if the inserted or filled bases match the truncated or deleted bases respectively. While figure 1c shows this for deletions, inserting ‘GC’ instead of deleting it results in the same sequenced barcode. Finally, we note that FREE divergence is not a metric—a mathematically precise term for distance—because edits outside the barcode window can lead to violation of the triangle inequality (Fig. 1c, Supplemental Materials). This requires us to use specialized code generation techniques that do not rely on the properties of a metric, and also underlies usage of the term divergence rather than distance throughout this work.

With FREE divergence defined, building an error correcting barcode list is conceptually equivalent to packing spheres in the space of possible barcodes (Fig. 2a). We set a barcode length *n* and call any DNA sequence of length *n* a word. For any word *B*, we call the set of all words *W* such that *FreeDiv(B, W)* ≤ *m* the *m-error decode sphere* of *B*, written as *DecodeSphere_m_(B),* or just *DecodeSphere(B)* if *m* is clear from context. Any observed DNA sequence within *DecodeSphere(B)* will by definition decode to (error-correct to) the center word *B* (Fig. 2a.). Then, an *m*-error correcting FREE code is simply any set of barcodes such that the *m*-error decode spheres of all barcodes are disjoint, i.e., no two decode spheres overlap. Any corrupted barcode with up to *m* errors is thus in the decode sphere of exactly one barcode and can be decoded (error-corrected) uniquely (Fig. 2a). Requiring disjoint decode spheres places a limit on the relationship between allowed *m*, the number of correctible errors, and *n*, the barcode length: to fit more than one non-overlapping decode sphere in the space requires that 2 *m* + 1 ≤ *n* (Supplemental Materials).

**Figure 2:**
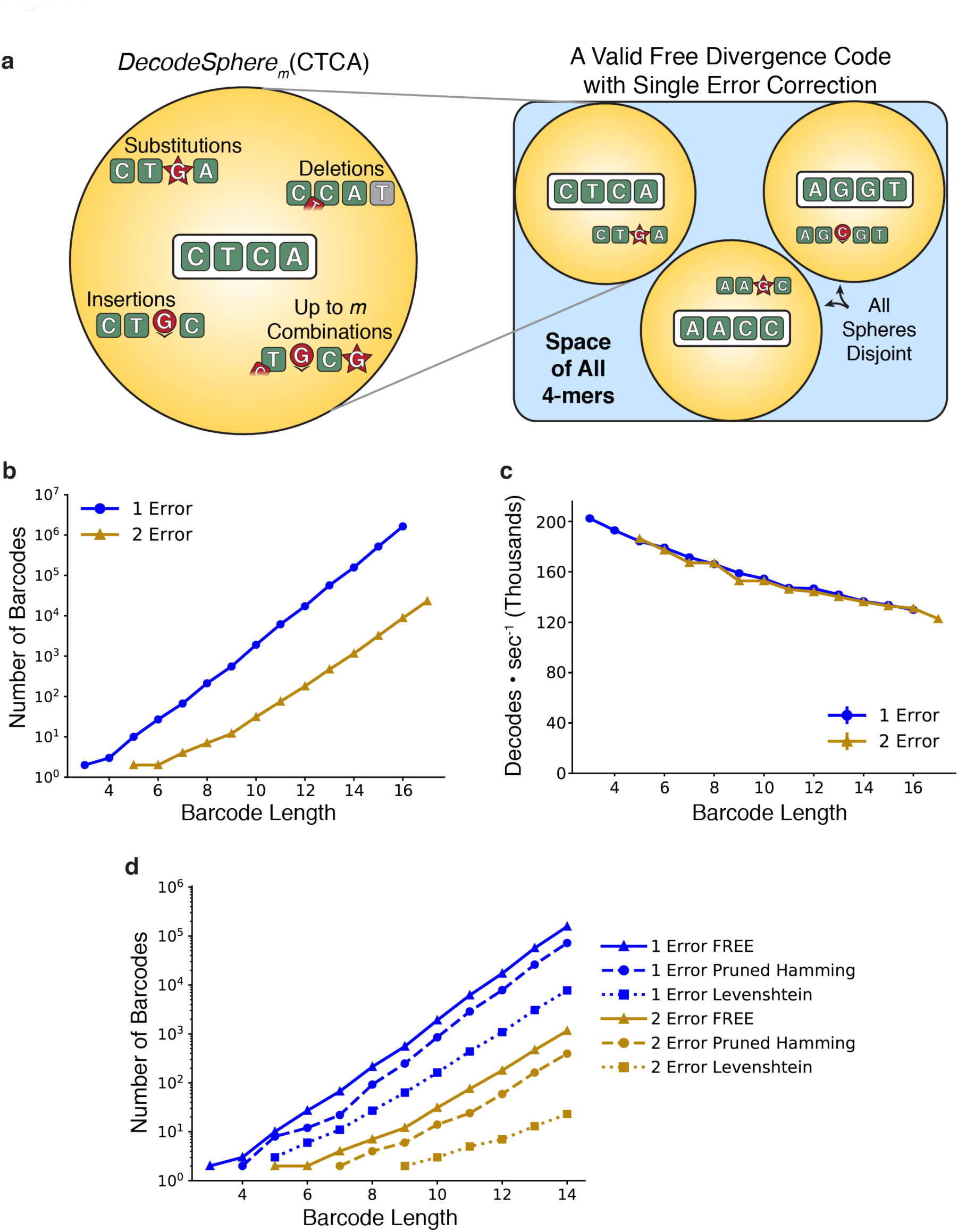
FREE barcode generation and decoding. **a.** Error-correcting barcode generation is a sphere packing problem. Around each accepted barcode *B* (e.g., “CTCA”), we reserve *DecodeSphere_m_(B)*, the set of all sequences within FREE divergence *m* of *B*. That is, the set of all sequences with any combination of up to *m* errors from *B*, followed by fill or truncation as necessary. Any set of disjoint decode spheres is a valid FREE code (right). **b.** The number of single-and double-error correction barcodes generated for a range of barcode lengths. **c.** The accompanying software decodes more than 120,000 barcodes per second for all barcode lengths considered here. **d.** Comparison of FREE barcode counts against pruned Hamming codes and Levenshtein codes. Hamming codes were pruned to remove members that did not decode FREE divergence errors, while Levenshtein codes were produced at double the error-correction levels for the same purpose. FREE codes produce more barcodes than either of the other methods for all barcode lengths.

### Efficient FREE barcode generation and decoding

A software library accompanying this manuscript efficiently generates FREE barcodes with a given total length and error-correction level. The generation algorithm is conceptually very simple: iterate through the space of n-mers alphabetically, find the decode sphere for each candidate barcode, and reserve barcodes when their decode spheres do not overlap the decode spheres of any previously reserved barcodes (Fig. 2a). This set of reserved barcodes by definition forms a valid FREE code. Additional algorithmic details make the process faster and more memory efficient (Methods). Adding valid code words in alphabetical order is a heuristic method previously observed to efficiently pack spheres24. Experimental synthesis and sequencing limitations are also incorporated during barcode selection. Candidate barcodes must have: (1) balanced GC content (40-60%); (2) no homopolymer triples (e.g., AAA); (3) no GGC (a known Illumina-based error motif^25^); and (4) no self-complementarity of >2 bases to reduce hairpin propensity. All of our software is available in the GitHub repository accompanying this manuscript (https://github.com/finkelsteinlab/freebarcodes).

The number of available error-correcting barcodes for a DNA sequence of length *n* will depend on the experimentally-required degree of error-correction (Fig. 2b). We generated libraries of single-error correcting codes up to a 16-nucleotide length, containing >1,600,000 barcodes. In addition, we generated more robust, double-error correcting codes up to a 17-nucleotide length with >23,000 unique members (Table S1). Barcodes correcting *m* errors require length at least 2*m* + 1 bp because otherwise all decode spheres overlap all other decode spheres (Supplemental Materials). Thus, the 1-error and 2-error correcting barcode libraries have minimum lengths of 3 bp and 5 bp respectively. All single-and double-error correcting barcode libraries shown in Figure 2b are included as supplemental data (Supp. File 1), and are available in the GitHub repository (https://github.com/finkelsteinlab/freebarcodes). The barcode decoding software runs in time proportional to the length of the barcodes but constant with respect to the number of barcodes in the library. Hence, 1-error and 2-error correcting codes decode at the same speed for a given barcode length even though the 1-error libraries contain many more barcodes (Fig. 2c). Even the slowest decodes considered here, the 17-mer double-error correction barcodes, decode at >120,000 barcodes • sec-1 on a desktop computer using a single processor.

### Comparison with current error-correcting DNA barcode strategies

Current state-of-the art error correcting DNA barcoding applications often use Hamming or Levenshtein error-correction strategies^20,23^. Hamming codes only correct substitutions, and are thus insufficient for any DNA barcode applications with indels^26^. However, they are linear codes, meaning the code words form a well-structured lattice in barcode space. We tested an alternative hypothesis that pruning these well-packed Hamming decode spheres to subsets with disjoint FreeDiv decode spheres could result in a more efficient packing—more barcodes for a given barcode length—than our alphabetical generation strategy. This was not, in fact, the case: FREE codes have about a factor of two more barcodes for a given length than our best pruning of Hamming codes (Fig. 2d).

Levenstein codes can be used directly (i.e., without pruning) because they account for indels, but must be used at 2-fold higher error correction for DNA barcode applications (Fig. 1b). We generated such over-corrected Levenshtein barcode sets in a manner similar to the FREE code generation strategy. This strategy produced even fewer barcodes than the pruned Hamming code sets. (Fig 2d, Methods). Sequence-Levenshtein codes attempted to solve the problems inherent in using Levenstein codes for DNA applications, but an error in the derivation of these codes often causes them to decode to the wrong barcode (Supplemental Materials)^27^. In sum, FREE codes offer a substantially larger number of usable barcodes for a given barcode length, when taking into consideration real-world errors such as deletions, insertions, and substitutions that are encountered during DNA sequencing and synthesis.

### Error Correction in Real and Simulated Data

We validated FREE barcodes generated in this study by both numerical simulation and experiment. Pooled oligonucleotide synthesis was used to produce a library of >8,000 oligos with double-error correcting barcodes at both ends (Fig. 3a). The barcodes were arranged such that each left barcode should only ever be observed on the same oligo with one specific right barcode sequence, and similarly for right barcodes. Hence, we were able to measure the rate of incorrectly decoding barcodes from observing unexpected left-right barcode pairs (Methods). We sequenced 1.4 million copies of this library on an Illumina MiSeq for an average coverage of 159x using the standard Illumina workflow.

**Figure 3:**
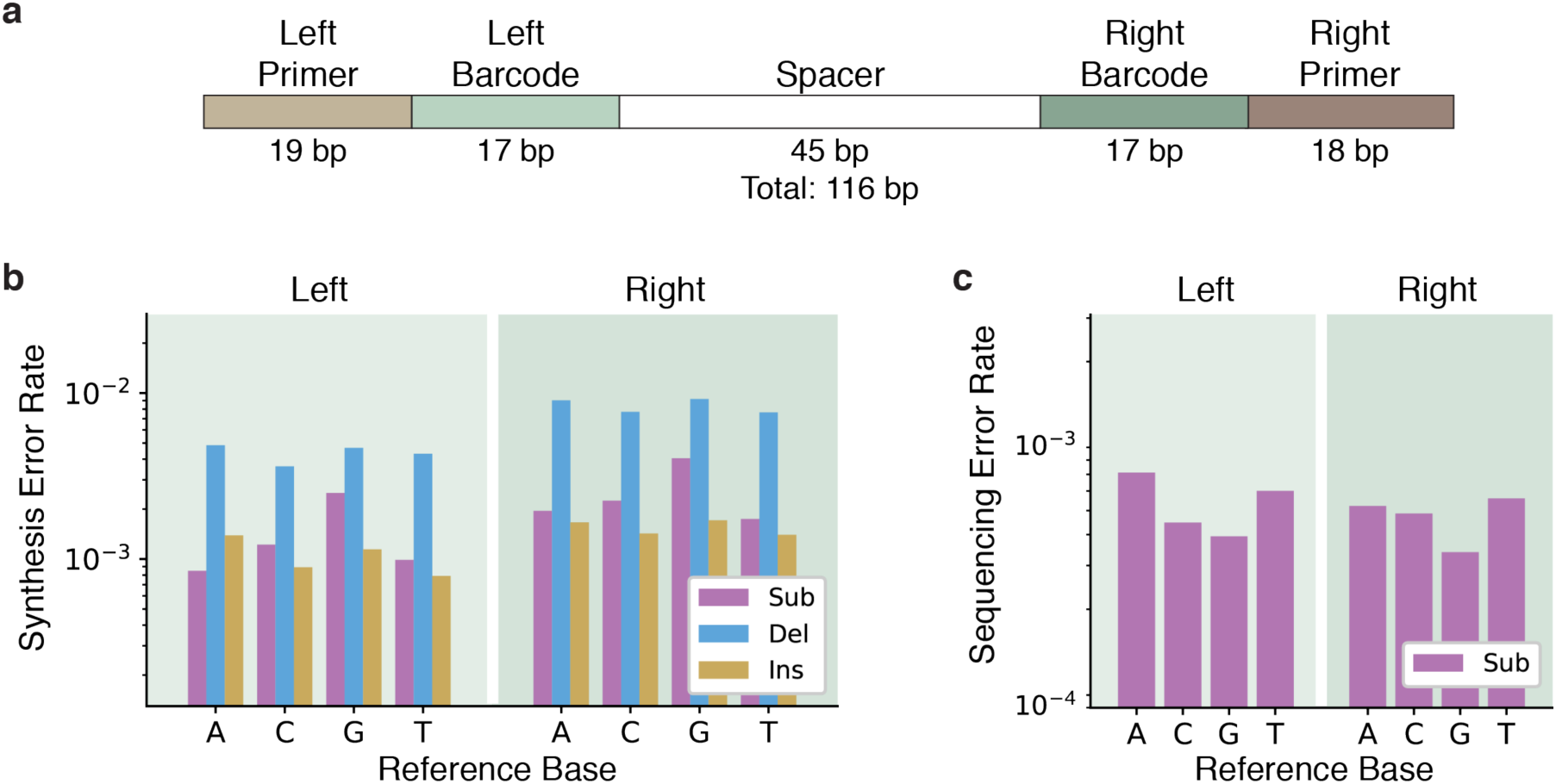
Experimental measurement of synthesis and sequencing error rates. **a.** Schematic of the DNA constructs used for barcode validation experiments. Each member in the synthetic library had a unique pair of left and right barcodes (green) drawn from a list of >8,000 17-nt FREE codes with double-error correction. By using the primer regions (brown) to distinguish the left and right ends from one another, we could determine whether the barcodes were correctly decoded (matching) or incorrectly decoded (mismatching). **b.** Synthesis error rates measured in this experiment, by intended reference base and error type—substitution (sub), deletion (del), and insertion (ins). **c.** Measured sequencing substitution error rates, by reference base. Insertions and deletions from Illumina sequencing are extremely rare and are omitted for clarity.

Full-length, paired-end Illumina sequencing was used to measure the background synthesis and sequencing error rates (Fig 3b-c). Using full-length paired-end reads permitted discrimination between synthesis and sequencing errors (Methods). Substitution, insertion, and deletion error rates from library amplification using Q5 polymerase have previously been reported to occur at rates less than 10^−5^, and thus are a negligible fraction of the measured synthesis errors^28^. Measured errors were dominated by single-base synthesis deletions, which occured at rates of approximately 1 in 200 bp and 1 in 100 bp in the left and right barcode regions respectively (Figs. 3b and S7). The two-fold difference in synthesis error rates between the two sides is consistent with statements from the manufacturer regarding their synthesis error rates^17^. Sequencing error rates are between 10^−4^ and 10^−3^, as advertised by Illumina (Fig 3c). In sum, experimental error rates are dominated by deletion errors. As Hamming codes are not designed to error-correct deletions in barcodes, they will perform very poorly in DNA-based experiments.

We compared the experimentally-determined error rates to simulations of the overall decoding error rate, i.e., the probability of incorrectly demultiplexing a barcode. Simulations were used to analyze the decode error rate for several error-correcting codes as a function of the per-base error rate, *p_err_* (Fig. 4). Simulations were performed in two different ways. First, we used a binomial model, which assumes independent and identically distributed errors at each base, to calculate the probability of observing more than 1- or 2-errors given per-base *p_err_*. Second, we directly simulated the errors directly using our decoding software: for a given per-base *p_err_*, we randomly select barcodes and add errors with probability *p_err_*. For simplicity, we model insertion, deletion, and substitution error rates of *p_err_* /3 with no correlation between individual errors within a given barcode. The corrupted barcodes are then decoded using our software and the fraction of incorrectly decoded barcodes is used as a measure of the decode error rate.

**Figure 4:**
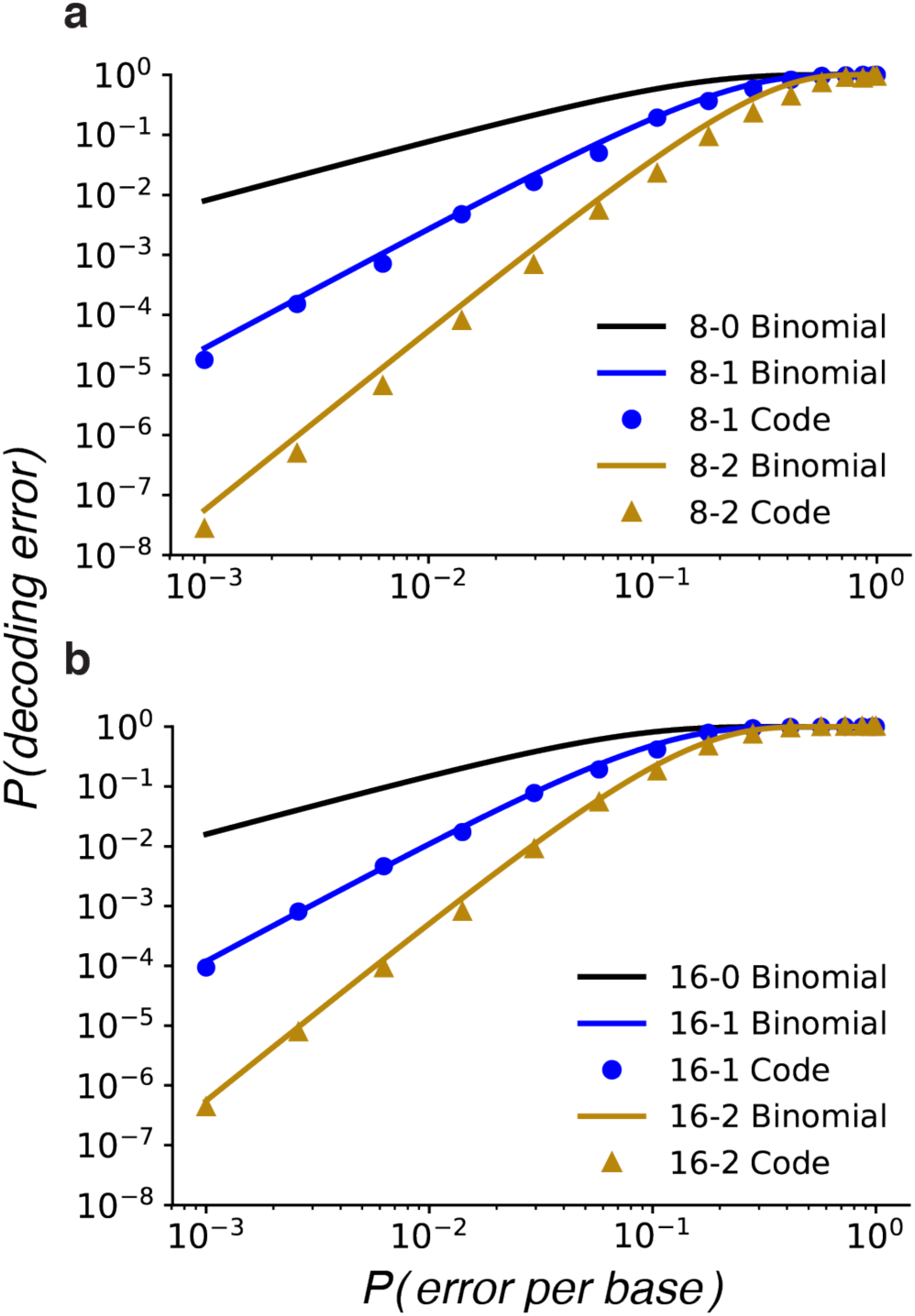
Decoding corrupted barcodes from simulated errors. Modeled and simulated decoding error rates given per-base error rate for length 8 (a) and length 16 (b) barcodes. Barcode sets are labeled according to length and number of errors corrected; for example, the 16-2 code is length 16 and corrects up to 2 errors. Solid lines show the error rate approximations using a binomial model. Circles and triangles show direct simulation error rates for single-and double-error correcting codes, respectively. Substitution, insertion, and deletion errors each have simulated error rate *P(error per base)*/3 for simplicity.

At experimentally-determined per-base error rates, *p_err_*, each increase in error correction level results in at least an order of magnitude improvement in the decoding error rate (Fig. 4). For example, our experimental data showed an overall per-base *p_err_* of approximately 10^−2^ (Fig. 3b-c). At this per-base error rate, the approximate uncorrected decode error rate (solid line) is 8% for length 8 barcodes and 15% for length 16 barcodes. Without error correction a best-case scenario would be that these errors could be successfully filtered out, representing a significant loss of data. In other scenarios, these data might be erroneously counted. For zero-, single-, and double-error correction length 8 barcodes, the approximate decode error rate decreases from 8% to 0.3% to 0.005%. For length 16 barcodes, the approximate decode error rate decreases from 15% to 1% to 0.05%. A more comprehensive comparison of the various barcode lists is given in figures S3-S5. The simulated results are consistently better than the binomial approximation because indels near the right end occasionally add the correct base and because insertions occasionally push other errors out of the barcode window (Fig. S2).

We validated FREE barcodes by measuring the decoding error rates for the experimental dataset described earlier (Fig. 5). For double-error correction, we used mismatches in barcode pairs to identify erroneously decoded barcodes (Methods). After corrections, we observe error rates of 0.29% and 0.46% for left and right barcodes respectively. We counted the 0- and 1-error correction rates shown in figure 5 by also counting the number of errors observed in each correctly decoded barcode. That is, 0-error correction decode error rates were calculated as the number of erroneously decoded barcodes plus the number of correctly decoded barcodes with 1 or 2 errors; 1-error correction errors were counted similarly. On the other hand, the theoretical model was calculated using the synthesis and sequencing error rates found in Fig. 3 to calculate the decode error probability of each barcode depending on its base composition, and then combined for an overall error rate (Supplemental Materials).

**Figure 5:**
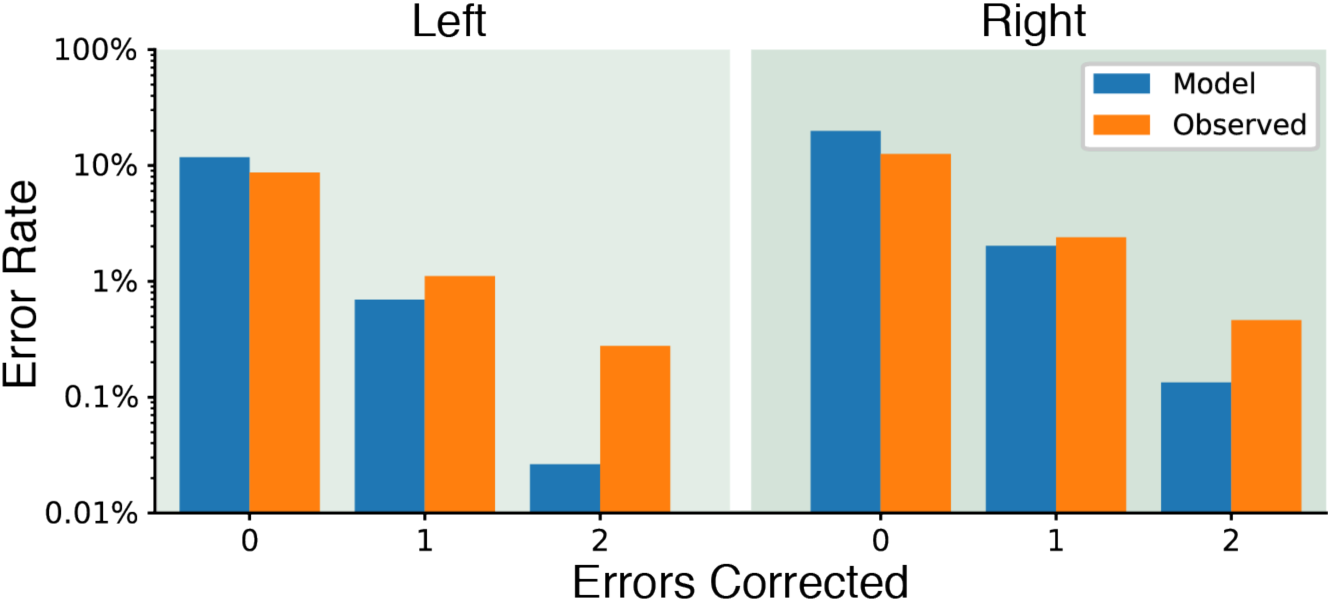
Decoding corrupted barcodes from experimental data. Observed decoding error rates compared with theoretical rates from the synthesis and sequencing error rates.

The experimentally-observed decoding error rates follow the same trend as the simulated errors: decode error rates decrease by approximately an order of magnitude with each additional error-correction level. We also observed that experimental error rates are higher than the theoretical error rate. This is explained by two observations. First, the theoretical model assumes independent errors at each position along the barcode. This assumption is not observed in the experimental data (Fig. S7). Second, the starting position of each barcode may not be defined exactly because the primer region can have errors. We are careful to identify the start of each barcode as precisely as possible (Supplemental Materials), but any errors in starting position appear as spurious insertions or deletions during decoding. Nonetheless, even though per-base errors are not independent, the overall order-of-magnitude decrease in decode errors per error-correction level is recapitulated in the experimental dataset.

### Combinatorially large barcode lists via concatenation

State-of-the-art high-throughput sequencing applications already require >106 unique barcodes^16^. We anticipate that improvements in high-density pooled oligo synthesis, along with the continuing reduction in sequencing costs, will continue to push the need for even larger error-correcting barcode sets. Below, we demonstrate that arbitrarily large barcode lists (>10^15^ unique members shown here) can be constructed from FREE barcodes by concatenating multiple FREE barcodes in a row.

As a demonstration, we concatenated two or three barcodes from the same starting list of sub-barcodes (Fig. 6). For the rest of this section we will refer to the original barcodes as *sub-barcodes*, while *barcode* will refer to the full length, concatenated barcode. Due to the possibility of insertions and deletions, the starting positions of the second and third sub-barcodes are only known approximately, and that approximation worsens as more sub-barcodes are added (Fig. 6a). Decoding the sub-barcodes sequentially from left-to-right is a strategy to account for this ambiguity. The left-most sub-barcode is decoded first, and then the decoded sub-barcode is used to find the starting position of the next sub-barcode. The error-correction level of each FREE sub-barcode remain the same, such that, for example, three concatenated double-error correction sub-barcodes can each correct up to two errors for a maximum total of six corrected errors if and only if the errors are evenly distributed, two per sub-barcode. Overall concatenated barcode decoding error rates are given by the probability of any decoding error in any sub-barcode or-barcodes. Concatenated barcode error rates are thus slightly higher than for the individual sub-barcodes (Fig. 6b). The decoding process is performed automatically using the software accompanying this paper.

**Figure 6:**
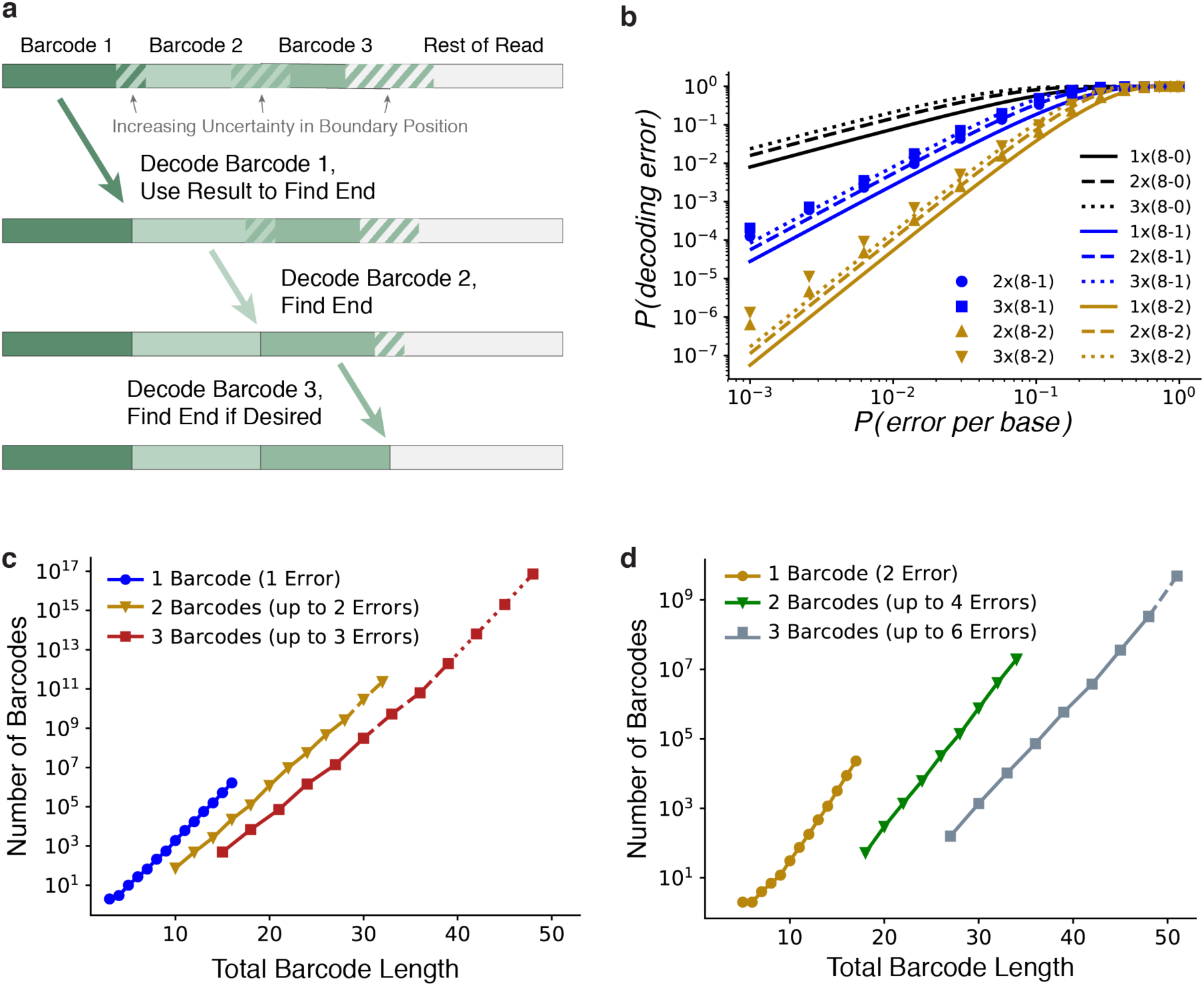
Combinatorial barcode libraries via concatenation of FREE barcodes. **a.** Concatenated barcodes can be decoded sequentially in a left-to-right order, even when the end position of each edited sub-barcode is not initially known. The decoded first FREE sub-barcode can be used to find the starting position of the next sub-barcode, and similarly for subsequent sub-barcodes. **b.** Concatenated barcode decoding error rates. Concatenated barcode labels use the following format: a 3x(16-1) barcode consists of three concatenated sub-barcodes, each of which is 16 bp long and can correct up to 1 error. Lines: binomial model. Points: direct simulation. **c, d.** Concatenating multiple barcodes combinatorially increases the numbers of effective FREE barcodes. Concatenated barcodes can correct the same number of errors per sub-barcode. When the errors are distributed evenly among the sub-barcodes, concatenated barcodes can correct a higher total number of errors than the individual sub-barcodes. (c) Concatenated single-error correcting barcodes. (d) Concatenated double-error correcting barcodes. Dashed lines: projected quantities calculated by sampling; dotted lines: log-linear projections.

Concatenating FREE barcodes results in combinatorially-large barcode sets that will be sufficient for even the most demanding high-throughput sequencing applications (Fig. 6). The concatenated barcodes were pruned to remain compatible with experimental constrains by removing DNA sequences that had triplet repeats of a single base or excess self-complementary (defined as any self-complementarity of any three or more bases). Even with these filters, we generated full lists of up to 10^10^ barcodes with concatenation of three single-error correcting codes (Fig. 6). Beyond that, where possible, the projected total barcode count was estimated via subsampling. When even that was limited by available hard drive space, the projected total was estimated via log-linear fit, which went above 10^15^ barcodes for 3 x (16 bp single-error) barcodes. Due to their size, we do not include these concatenated barcode sets explicitly with this paper. They can be generated on demand using the included software package and single barcode lists. In sum, concatenating FREE codes produces a rapid and efficient strategy for further increasing the size of error-correcting barcode lists for pooled high-throughput sequencing experiments.

## DISCUSSION

Here, we described the design and experimental validation of Filled/truncated Right End Edit (FREE) error-correcting DNA barcodes capable of correcting substitution, insertion, and deletion errors, even when the corrupted length of the barcode is unknown. We generated lists of FREE Divergence error correcting barcodes and provided software on GitHub for user-friendly generation and decoding of these DNA barcodes for real-world applications.

Most high-throughput DNA sequencing applications require PCR-based amplification or reverse transcription (in the case of RNA) of the input nucleic acid libraries. The polymerase and reverse transcriptase enzymes used during library preparation perform best on libraries that avoid stable secondary structures and self-complimentary regions. To improve the utility of our codes for such demanding applications, we used UNAFold to calculate the melting temperature of hairpins for the FREE barcodes included with this paper29. This information will allow users to prune out barcode sequences with a propensity to form stable hairpins in their specific experimental conditions (Fig. S8). Such experimental considerations will further increase the utility of FREE codes for demanding high-throughput sequencing applications.

In validating the FREE barcodes, we measured the types and frequency of errors that are introduced during massively-parallel oligo synthesis and Illumina-based high-throughput sequencing. We observed that deletions during synthesis were the most frequent sources of error (~1 per 100 nucleotides), followed by substitutions and insertions (~1 per 1000 nucleotides). These experimentally measured error frequencies were used to simulate and experimentally measure the decoding quality of FREE codes. Even though the observed decoding error rates do not follow a model that assumes independent errors at each base, we still obtain exponential improvement of the final decoding error rate with codes that correct for increasing numbers of errors. Importantly, the error-correcting decode software runs fast enough to handle the massive data sets involved in modern high-throughput sequencing applications, decoding hundreds of thousands of barcodes per second on a single processor for all barcode lists considered.

While we have here focused exclusively on filled/truncated *right* end edit (FREE) codes prepended to the start of sequenced DNA reads, the current work applies equally to their natural mirrored counterpart, filled/truncated *left* end edit (FLEE) codes. This would be required for applications where the barcode appears at the end of each sequenced read rather than the beginning. In fact, the same codes can be used by simply taking the reverse complement of FREE codes before synthesis and again before decoding. Hence, FREE barcodes can be used equally well on the 5’ or 3’ end of pooled samples, as long as the orientation is chosen appropriately.

FREE barcodes are a powerful tool to correct DNA barcode errors, reducing measurement errors in modern, high-throughput experiments. We anticipate that the use of FREE barcodes will improve these assays in three key ways: (1) helping avoid spurious results; (2) decreasing the amount of discarded data; and (3) increasing experimental signal-to-noise ratios. Decreasing spurious results and discarded data are important for any experiment involving DNA barcodes, but we are most excited by the new possibilities available with increased signal-to-noise ratios. The power to decrease error rates from 15% to 0.05%, as in Fig. 4b, could open the door for entirely new assay designs. We anticipate that FREE barcodes will be broadly useful for the ever-growing set of pooled high-throughput sequencing experiments in cell and molecular biology, protein engineering, and drug discovery.

## METHODS

### Definitions and Numerical Representation of DNA

For any barcode system, the word length, *n,* is given. Any DNA sequence of length *n* is a *word*, and any word observed in the data is an *observed word*.

We represent strings of DNA as base-4 numbers where A, C, G, and T correspond to 0, 1, 2, and 3 respectively. So, for example,

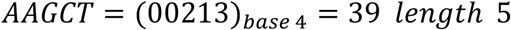

Here 39 is the *word number* and 5 is the *word length.* Note that the word length is required to uniquely convert numbers to DNA to account for leading A’s. For example, the word number from the example above, 39, with word length 3 is simply GCT. For word length *n*, the largest valid word number is 4^*n*^ – 1.

For an *m*-error correcting code we define a *decode sphere* around a barcode *B* to be the set of all words with FreeDiv less than or equal to *m,* and we define an *encode sphere* to be the set of all words of FreeDiv less than or equal to 2*m.* We write these as *DecodeSphere(B)* and *EncodeSphere(B)*.

### Barcode Generation

FREE barcode sets are generated with a modified lexicographic code generation method. Lexicographic code generation consists of marching through all words lexicographically, alphabetically in this case, and adding new words to the list of barcodes whenever they are sufficiently far from all previous barcodes^30^. For Hamming codes, lexicographic codes are linear30, and more generally, lexicographic code generation has been shown to have relatively good sphere packing efficiency^24^. The first FREE modification to the procedure is to enforce the following sequencing and synthesis properties:

- Balanced GC content (40-60%)
- No homopolymer triples (e.g., TTT)
- No triplet self-complementarity
- No GGC (Illumina error motif^25^)

For speed we iterate over these potential barcodes via recursive base addition: given a barcode prefix *P*, we add the next base only if it does not violate any of the above. We thereby skip large recursive subtrees in which all words violate one of the above conditions.

For an *m-*error correcting code, the only requirement is that the decode spheres of all barcodes are disjoint. Because FREE divergence is not a metric, standard metric-based code generation methods cannot be used. Instead, we accomplish this directly with a sphere iterator (Supplemental Materials). For every accepted barcode *B*, we iterate over *DecodeSphere(B)* and reserve all words therein as mapping to *B.* And for any potential new barcode *P*, we first verify no words in *DecodeSphere(P)* are reserved before accepting it as a new barcode.

This algorithm would be very slow because most decode sphere tests would run into reserved words and fail to add new barcodes. One further observation makes this process tractable. Given a barcode B and a proposed new barcode *W,* if *FreeDiv(B, W) ≤ 2m*, that is, if *W* is in *EncodeSphere(B)*, then *DecodeSphere(W)* and *DecodeSphere(B)* overlap and *W* is not a valid new barcode (Supplemental Materials). This implies the following algorithm: generate the code by lexicographically iterating over words while looking for new barcodes to add to the code. For each accepted new barcode *B*, we color any uncolored words in *EncodeSphere(B)* black, and then we color all words in *DecodeSphere(B)* red. Restricting encode sphere coloring to previously uncolored words avoids overwriting the decode spheres of all previous barcodes. All black-and red-colored words are guaranteed to not be valid barcodes, so addition of new barcodes is restricted to uncolored words. For an uncolored proposed new barcode *W*, *DecodeSphere(W)* is checked for red words. If no red words are found, *W* is added as a new barcode.

The coloring of barcodes, decode spheres, and encode spheres is accomplished by having an array of 4^*n*^ integers valued 0, 1, or 2: 0 for uncolored, 1 for black, and 2 for red. The location of each integer in memory itself represents the word, via the numerical representation of DNA given above. This is both memory and speed efficient. Memory efficiency is important, as it is a limiting resource for this method. The memory required for barcode generation is 4^k^ bytes, which for this paper was up to 16Gb of random access memory (RAM).

### Barcode Decoding

The decoding process builds the code book and looks up decoded words directly. We do this in a memory efficient fashion as follows. For each barcode in a list, the *barcode index* is defined as the index of that barcode within the list of barcodes. We again reserve a space of 4^*k*^ integers to represent the code space. For each barcode *B*, we store the barcode index of *B* at every word of *DecodeSphere(B).* We store barcode indices rather than barcode numbers because barcode indices require fewer bits per word. The memory required for barcode decoding is (1, 2, or 4) × 4^*n*^ bytes, requiring 1, 2, or 4 bytes to store each barcode index. For this paper, the maximum memory used for barcode decoding was 32Gb of RAM.

### Barcode Pruning

Specific barcode lists from literature or elsewhere may sometimes be required for a given experiment, but require pruning to find a subset with error-correction. We accomplish barcode pruning via the same strategy as barcode generation, but only considering the input set of barcodes as potential new barcodes. This pruning method was also used to prune the linear Hamming codes.

### Simulation of Errors

To test the error-correcting capacity of FREE barcodes, we wrote error-simulating code which adds a given number of substitutions, insertions, deletions, or all three randomly distributed. We used this to verify the correctness of each of the FREE *m-*error correcting codes by randomly selecting barcodes, adding *m* errors, and verifying that the decoded word matches the expected word. We used the same code for generating Figure 4 by randomly choosing the number of errors from a binomial distribution with probability of error *p_err_.*

### Levenshtein Barcodes

Levenshtein barcodes were generated lexicographically using the standard technique of code generation with a metric. Briefly, for desired barcode length *n* and number of correctable errors *e,* we walk through the space of *n*-mers lexicographically adding any new word if it: (a) satisfies the same sequencing and synthesis properties as above, and (b) is Levenshtein distance at least 2*e*+1 from any previously accepted barcode.

### Pruned Linear Hamming Barcodes

We generated Hamming barcode lists using linearity in base-4 (Supplemental Materials). Briefly, a Hamming code of length *n* with *k < n* “message bits” can be generated by all linear combinations of *k* basis vectors of length *n* which are chosen to enforce the error-correction properties required. These codes were then filtered according to the FREE sequencing and synthesis property requirements and pruned as described above to form valid FREE codes.

### Experimental Synthesis, Sequencing, and Decoding Error Rates

Oligonucleotide pools were designed as in Figure 3a, with primers and barcodes on each end and a spacer in the middle (116 bp total length). To test the FREE method, 8,634 barcodes of length 17 and double-error correction were used in 8,634 unique pairs. Oligos were synthesized (CustomArray), and the oligo pool was amplified for twenty cycles with Q5 polymerase (NEB) and sequenced on an Illumina MiSeq machine with 2×150 bp paired-end reads. Maximum likelihood sequences were inferred using both reads.

The left and right primer sequences were used to determine both the read orientation and the starting position of each barcode (Supplemental Materials). Each barcode was then decoded using the FREE decoding software. Matching barcodes identified correctly decoded barcodes, while mismatching barcodes indicated an error. The FREE method was powerful enough to reveal a surprising and unrelated source of error: the creation of oligo chimeras, sequences with the left part of one oligo and right part of another, which we then also accounted for (Supplemental Materials).

Once each oligo had been identified from its barcodes, the observed sequence was aligned with the reference sequence. At each base where the two reads agreed with each other but not with the reference sequence we counted a synthesis error, at each base where the reads disagreed and one read matched the reference sequence we counted a sequencing error, and at each base where the reads disagreed and neither matched the reference sequence we counted a synthesis and a sequencing error.

Observed synthesis and sequencing error rates for each reference base were used to find theoretical decoding error rates for each barcode given its base composition. These were then used to estimate overall expected error rate (Supplemental Materials).

## ACKNOWLEDGMENTS

This work was supported by a College of Natural Sciences Catalyst award, the Welch Foundation (F-1808 to I.J.F.), and the National Institutes of Health (R01 GM120554 & GM124141 to I.J.F., F32 AG053051 to S.K.J.). We thank James Rybarski, Andrea Hawkins-Daarud, Jeffrey Hussmann, Prakash Mohan, Alexander Boulgakov, and Kevin Drew for useful feedback throughout the project.

## AUTHOR CONTRIBUTIONS

J.A.H., I.J.F, and W.H.P. designed the research. J.A.H. wrote the software and analyzed the experimental data. J.A.H. and S.K.J. prepared the DNA for sequencing. J.A.H., I.J.F., and W.H.P. wrote the paper. All authors commented on the manuscript.

